# Echo State Networks Ensemble for SSVEP Dynamical Online Detection

**DOI:** 10.1101/268581

**Authors:** D Ibanez-Soria, A. Soria-Frisch, J Garcia-Ojalvo, G Ruffini

## Abstract

**Background:** Recent years have witnessed an increased interest in the use of steady state visual evoked potentials (SSVEPs) in brain computer interfaces (BCI), SSVEP is considered a stationary brain process that appears when gazing at a stimulation light source.

**New Methods:** The complex nature of brain processes advocates for non-linear EEG analysis techniques. In this work we explore the use of an Echo State Networks (ESN) based architecture for dynamical SSVEP detection.

**Results:** When simulating a 6-degrees of freedom BCI system, an information transfer rate of 49bits/min was achieved. Detection accuracy proved to be similar for observation windows ranging from 0.5 to 4 seconds.

**Comparison with existing methods:** SSVEP detection performance has been compared to standard canonical correlation analysis (CCA). CCA achieved a maximum information transfer rate of 21 bits/minute. In this case detection accuracy increased along with the observation window length

**Conclusions:** According to here presented results ESN outperforms standard canonical correlation and has proved to require shorter observation time windows. However ESN and CCA approaches delivered diverse classification accuracies at subject level for various stimulation frequencies, proving to be complementary methods. A possible explanation of these results may be the occurrence of evoked responses of different nature, which are then detected by different approaches. While reservoir computing methods are able to detect complex dynamical patterns and/or complex synchronization among EEG channels, CCA exclusively captures stationary patterns. Therefore, the ESN-based approach may be used to extend the definition of steady-state response, considered so far a stationary process.

**Highlights:**

- We present a novel SSVEP dynamical detection approach based on ESN.
- This is the first time ESNs are applied to SSVEP based BCI systems.
- We provide experimental validation of proposed methodology.
- Experimental results indicate non-stationarity in SSVEP patterns.

## I. Introduction

A Brain Computer Interface (BCI) provides a direct communication pathway connecting the brain to a computer or other external device. BCIs do not rely on the brain’s normal action pathways through peripheral nerves and muscles, making them an ideal technology for systems assisting or repairing human cognitive or sensory-motor functions [1]. Different modalities for the realization of BCI exist; the most commonly used ones being motor imagery, P300, and Steady State Visual Evoked Potentials (SSVEP) [2]. SSVEP-based BCIs offer two main advantages [3]: i) they have a larger information transfer rate, and ii) they require a shorter calibration time.

Many SSVEP-based BCI systems rely on a protocol where the subject decides voluntarily when to interact with the BCI application. When the system detects the user’s intention to perform an action, the visual stimulation is presented during a short time period. A typical SSVEP-based BCI system with *N*_*s*_ degrees of freedom employs *N*_*s*_ independent light sources flickering at different frequencies. Each light source is associated then to a particular action of the BCI system. Therefore, when the user wants the system to perform a specific action, he\she shall gaze at its associated light source. Detection methodologies aim at determining immediately after the short-time stimulation period which of the stimulation frequencies in the BCI set-up elicited a visual evoked response.

Steady-state visual evoked potentials, which appear when a person gazes at a flickering light source, are measured by electroencephalography (EEG). The flickering frequency of the light source can range from 1 to 100 Hz [5]. SSVEPs appear as oscillatory components in the user’s EEG matching the stimulation frequency and its harmonics, they are mainly observed in the primary visual cortex, and are characterized by an energy increase that is phase-locked with the visual stimulus [5]. The SSVEP response is a subject- and stimulation-dependent phenomenon determined by the stimulation frequency, its intensity, color, and duty cycle [2].

Figure 1 depicts the average spectral response over various stimulation trials for one subject while gazing at a 12 Hz stimulus along with the average spectral response of sequences with no visual stimulation. An energy increase at the stimulation frequency (12Hz) and its second harmonic (24 Hz) can be clearly observed during visual stimulation. While the detection of such a response seems simple when using the average response over several stimulation trials, it becomes very difficult when attempting it over single short-time stimulation trial periods. The objective of this paper is to address this issue.

**Fig. 1.**
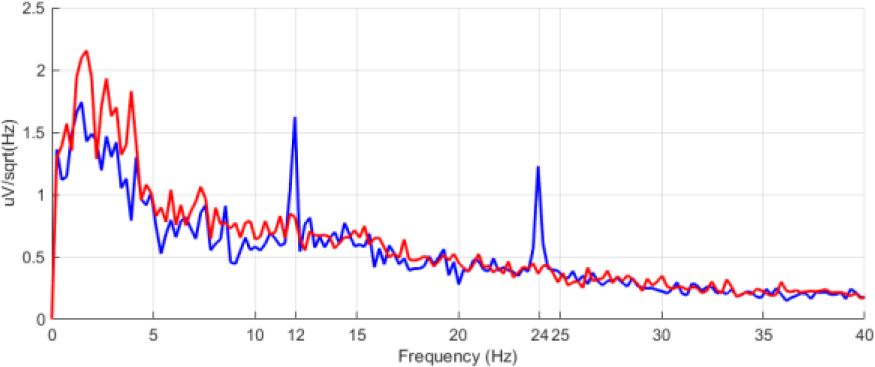
Average spectral response in a particular subject when gazing at a 12 Hz visual stimulation (blue) and in sequences of no visual stimulation (red).

The brain is a complex, dynamical system that generates non-stationary EEG patterns [6]. This complexity limits the efficacy of non-dynamical feature extraction and classification methods. In this context, it is worth mentioning that the study of transient responses of complex dynamical systems has recently led to the proposal of a new information-processing approach, known as Reservoir Computing (RC). This paradigm opens a wide variety of possibilities for EEG analysis in general and for real-time applications, such as BCI in particular. Although RC techniques have been applied with excellent results to motor imagery [7], they have never been used for SSVEP detection before (to our best knowledge). This is probably due to the assumed stationary nature of the SSVEP response. In this work we aim to exploit the capabilities of reservoir computing for online SSVEP detection. The performance of the proposed RC approach is compared to standard canonical correlation analysis (CCA), which has shown better performance than traditional fast Fourier transform-based spectrum estimation methods [8].

The paper is structured as follows. In section 2 we provide an overview of reservoir computing. We present the new methodology based on RC in section 3. In section 4 we describe our experimental protocol, and in section 5 we provide results for the two approaches under comparison. We conclude with a discussion in section 6.

## II. Reservoir Computing

Artificial neural networks have been extensively used for the analysis of stationary problems in computational intelligence. These architectures are well understood due to their feed-forward structure and non-dynamical nature. It is in general not possible to detect temporal dynamics using feed-forward structures. A possible temporal generalization strategy is to add recurrent connections, which allow the system to encode time-dependent information by providing the network with fading memory and transforming it into a complex system [9]. In contrast with feed-forward networks, neural networks whose activation is fed unidirectionally from input to output, recurrent neural networks (RNNs) present at least one cyclic path of synaptic connections, implementing a nonlinear dynamical system [10]. Dynamical systems are commonly used to model non-stationary physical phenomena. These models are extensively used in a wide variety of fields including finance [11], economics [12] and physiology [13].

Since the early 1980s, a wide variety of approaches for adaptive learning in networks with recurrent connections have been proposed [14]. Training RNNs has traditionally been more complex and computationally more expensive than training feedforward neural networks. Additionally, cyclic connections can provoke nonlinear bifurcations leading to drastic changes in its behavior [15]. Echo State Networks (ESN) [16] and Liquid State Machines [17] together constitute a new approach towards training and applying Recurrent Neural Networks, grouped under what came to be known as reservoir computing (RC). RC is based on the principle that supervised adaptation of all interconnection weights in RNNs is not necessary: training a supervised readout from the reservoir is sufficient to obtain excellent performance in many tasks [18]. This approach has certain analogies with kernel methods in ML, with the reservoir performing a nonlinear high-dimensional projection of the input signal for discriminating samples that are not linearly separable in the original space. At the same time, the so-called dynamical reservoir, which is formed by a fully connected network of varying number of nodes, serves as a memory providing the temporal context [19].

The global structure of an RNN is depicted in Fig. 2. Following the nomenclature and model adopted by Jaeger in [10] we consider a network of *K* input units, N internal units, and L output units. Input, internal and output connection weights are defined respectively by the connection weight matrices *W*^*in*^ (*N* × *K*), *W* (*N* × *N*) and *W*^*out*^ (*L* × (*K* + *N*)). RNNs are characterized for having cyclic paths of synaptic connections within *W*. Additionally, a back-projection weight matrix *w*^*back*^ (*N* × *L*) can be considered. In ESN-based supervised training the random input (*W*^*in*^), internal (*W*) and back-propagation weights (*W*^*back*^) matrices form the dynamical reservoir (DR). A DR is an echo state network if it presents the echo state property. This property states that the current state of a network, which is running for an infinite time, is uniquely determined by the history of the input and the teacher-forced output (i.e. the initial state of the RNN does not matter, since it is forgotten). The echo state property has proved to be linked with the characteristics of the reservoir, with the input signals and with the input and back-propagation weights [20,21,22]. The weight matrix is usually characterized by its spectral radius, defined as the largest absolute eigenvalue of the weight matrix. It is closely connected with the intrinsic dynamical timescale of the reservoir, and is therefore a key ESN training parameter. A small spectral radius leads to a faster RNN response. In most practical applications, a spectral radius below unity ensures the echo state property [23]. Other key training parameters in RC are the input scaling and the model size [23]. The model size is defined by the number of internal units N. Generally, a larger DR can learn more complex dynamics, or a given dynamics with greater accuracy. It is very important however to be aware of the possible over-fitting if a large number of internal units is chosen, which would lead to poor generalization [10]. The input scaling determines the degree of nonlinearity in the reservoir responses. Tasks close to linear require small input scaling factors, while highly nonlinear tasks demand larger input scaling values.

**Fig. 2.**
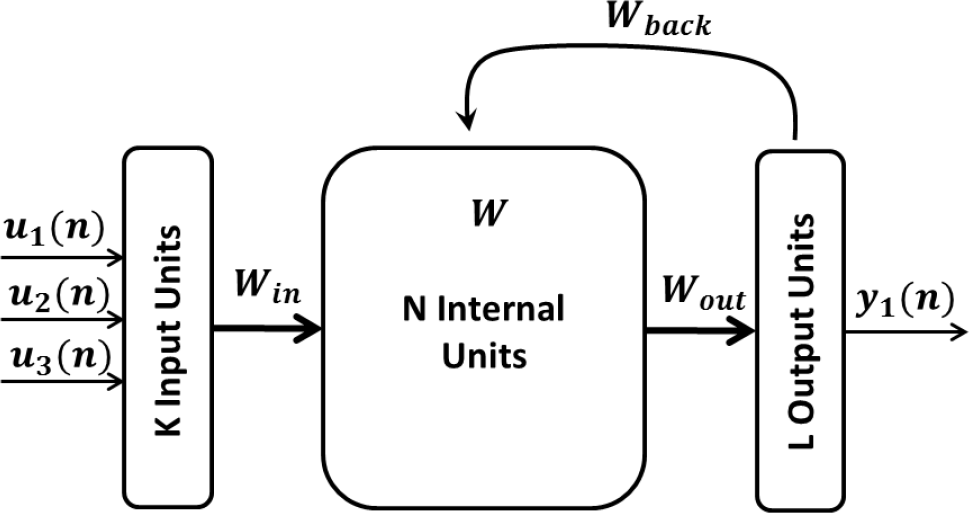
Recurrent Neural Network structure configured as the one proposed for SSVEP detection, 3 input units and 1 output unit (K =3 and L=1).

## III. Methods

### A. Canonical Correlation Analysis

Canonical correlation analysis (CCA) [24] is a multivariable calibration-less statistical method to calculate the maximal correlation between two multi-channel signals. CCA is widely used in statistical analysis and information mining [25, 26]. SSVEP-based BCI systems have largely used CCA-based methods in recent years due to its excellent accuracy and information transfer rate [27, 28]. Given two multidimensional random variables *X*, *Y* and their linear transformation *x*̃ = *w*^T^*X* and *y*̃ = *v*^T^*Y*, CCA finds the weight vectors *w* and *v* that maximize the correlation between *x*̃ and *y*̃. Canonical correlation therefore seeks a pair of linear transformations for *X* and *Y* such that when the multidimensional variables are transformed, the corresponding coordinates are maximally correlated [29]:

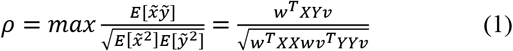

The SSVEP response is characterized by oscillations in the visual cortex matching the stimulation frequency and its harmonics. The performance of a given stimulation frequency *f*_*i*_ is evaluated by computing the canonical correlation between the EEG sequence under evaluation (*X*) and a reference signal (*Y*), constructed as a set of sine-cosine series at the stimulation frequency and its *N*_*h*_ harmonics of duration equal to that of the EEG sequence.

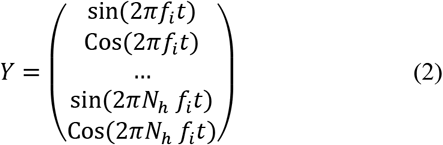

The maximal canonical correlation (*ρ*_*i*_) is calculated for all *N* stimulation frequencies (*f*_*i*_) being tested. In standard CCA the stimulation frequency delivering the largest canonical correlation (*ρ*) is selected as responsible of eliciting the visual response [30].

### B. Reservoir-Computing Ensemble SSVEP Detection

In this section, we propose a novel approach for extraction and classification of SSVEP features, based on an ensemble of as many echo state networks as degrees of freedom the SSVEP-based BCI application has. Each ESN is trained to detect the elicited response to a particular stimulation frequency. The proposed approach is able to detect linear and nonlinear patterns in the EEG response, boosting the capabilities of state-of-the-art stationary detection methodologies.

#### 1) Temporal SSVEP Feature Extraction

SSVEP EEG temporal components are calculated for each of the *N_e_* electrode signals (which are denoted as *x*_*i*_(*n*), *i* = 1,2,3 … *N*_*e*_). The SSVEP temporal response at *f*_*i*_ for harmonic *K* is computed by filtering the raw EEG using a band-pass FIR filter (1 Hz bandwidth) with its central frequency at *K* · *f*_*i*_ The ensemble response of every harmonic under evaluation is computed by adding the calculated temporal response of each harmonic, obtaining the 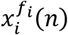 vector.

#### 2) ESN Construction

ESNs (one per stimulation frequency) have been configured to have one output node, and as many input nodes as EEG electrodes measure the visual evoked response (*N*_*e*_). As will be explained in following sections, the optimal number of internal units, spectral radius, and input scaling factor are calculated using three-fold cross validation exhaustive search.

The temporal decomposition of SSVEP components coming from each electrode 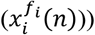 feeds the *N*_*e*_ input nodes of the ESN targeting detection of *f*_i_. The proposed ESN detection and classification methodology is applied to an EEG recording acquired during the interleaving of *N*_*s*_ non-stimulation periods followed by *N*_*s*_ stimulation ones, where the visual stimulation is presented at *f*_*i*_. The function of the ESN is to discriminate visual stimulation periods at *f*_*i*_ from non-stimulation periods or stimulation periods at other frequencies. For this two-class classification problem, during ESN training, the network outputs 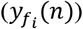 of samples corresponding to stimulation periods at *f*_*i*_ are set to 1, while the output of samples corresponding to non-stimulation periods are set to −1. Therefore, during the ESN recall the associated ESN output is maximized during the visual stimulation period.

**Fig. 3.**
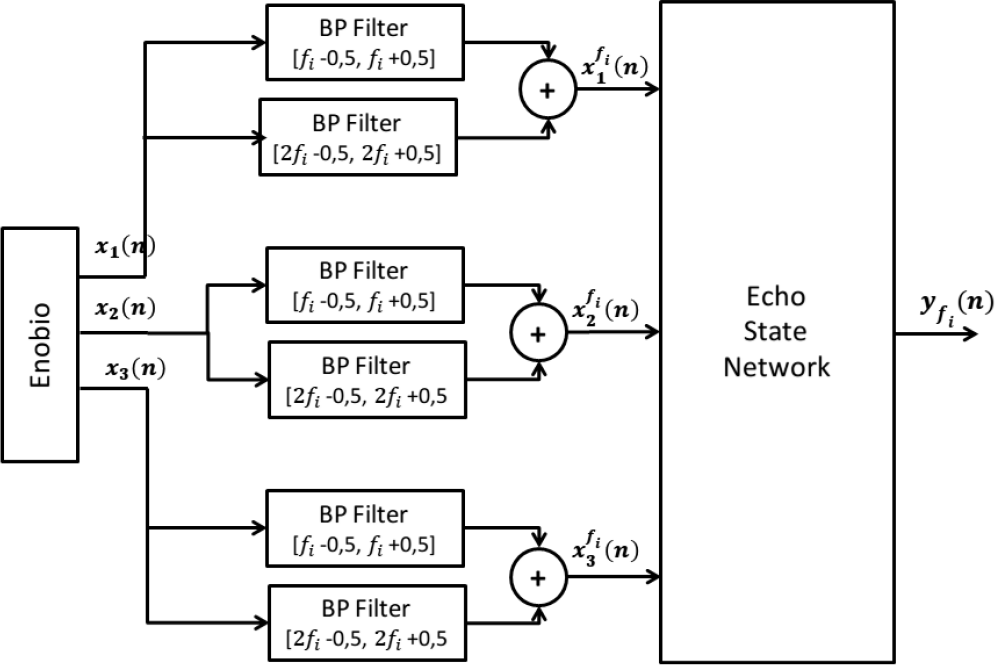
Proposed RC-based SSVEP Feature Extraction Architecture for the detection of the stimulation frequency ***f***_***i***_.

#### 3) Stimulation Frequency Detection

SSVEP frequency detection aims at determining which stimulation frequency among the ones under evaluation (*f*_*i*_, with *i* = 1,2 …*N*) elicited the visual response during a stimulation trial. To achieve this, the RC-based architecture has been designed as an ensemble of *N* ESNs, where each ESN has been trained for each particular stimulation frequency *f*_i_. The SSVEP response at each frequency 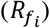 is assessed as the difference between the averaged ESN output 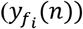 during the stimulation observation window and a baseline sequence prior the stimulation. As the ESN output is maximized during stimulation periods at the trained stimulation frequency, the stimulation frequency with maximal 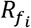 will be selected as responsible of eliciting the visual response.

## IV. Experimental Method

Five Caucasian male subjects S1 to S5 with average age 33.6 years participated in six recording sessions each, where oscillatory visual stimuli were presented at six different frequencies *f*_*i*_: 12, 14, 16, 18, 20 and 22 Hz. Although a strong SSVEP response can be obtained for stimulation frequencies in the range 5-20 Hz [31], stimulation frequencies below 12 Hz were discarded because the subjects found them uncomfortable. The visual stimuli were presented using stimulation sources consisting of an array of flickering light emitting diodes (LEDs) through a diffusing panel of 100 squared centimeters, developed specifically for this study. Stimulation sources integrated a communication layer through which their stimulation frequency, duty cycle, luminosity and color could be configured. In this study, the LED current was modulated in the form of a squared 50% duty-cycle excitation, with white color and maximum luminosity for every stimulation frequency.

Each session consisted of one recording per stimulation frequency. In each recording, *N*_*s*_ = 15 stimulation trials (duration randomly ranging from 4 to 5 seconds), where the visual stimulus was presented, were followed by the same number of non-stimulation trials (duration randomly ranging from 5 to 8 seconds) with no visual stimulation. Visual stimuli were presented using two stimulation sources next to each other. Higher number of simultaneous stimulation sources, which would reproduce for instance a BCI system of 6 degrees of freedom, were not used because of limitations in the available hardware and stimulus-presentation platform. In our case, one stimulation source was placed on the right of the subject, presenting the stimulation frequency under evaluation, while the one placed on the left presented a frequency randomly selected among the other frequencies used in the experiment. Stimulation sources were separated by approximately 25 cm. The user was comfortably seated at one-meter distance from the stimulation sources and was instructed to look at the stimulation source placed on his right when hearing a beep sound (played one second before the stimulation started). EEG was acquired using an Enobio^®^ recording system at a sampling rate of 250 samples/second from three channels placed in O1, Oz and O2, according to the 10-20 system [32], with the electrical reference placed in the right ear-lobe. Background ambient light remained homogeneous throughout all experimental sessions. The AsTeRICS [33] platform was used to record the EEG streaming data, control the stimulation panels, and trigger the recording.

## V. Performance Evaluation and Results

The goal of the performance evaluation is to determine whether RC-ESNs are a suitable tool to be used in practical SSVEP-based BCI applications. To do so its performance is compared to standard canonical correlation detection. Although, as mentioned above, all six stimulation frequencies being tested were not simultaneously presented due to hardware limitations, a BCI system with six degrees of freedom is simulated. In operational conditions the detection methodologies will aim to determine which of the 6 frequencies under evaluation (12, 14, 16, 18, 20 and 22 Hz) is responsible for eliciting the evoked potential after the stimulation trial. Self-paced BCIs require the system to prompt the user for a response and therefore ignore unexpected user input. The performance of such systems is usually measured using the information-transfer rate *B* [34], which quantifies (in bits per minute) the amount of information reliably received by the system [35]. Denoting the application speed in trials/second by *V*, the classifier accuracy by *P* and the number of degrees of freedom of the BCI system by *N*, the information transfer rate is calculated as:

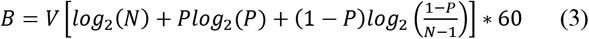

### 1) Canonical Correlation Analysis

Recordings have been band-pass filtered using a finite impulse response filter with 250 coefficients and high and low cut-off frequencies set to 1 and 45 Hz, respectively. The filtering aims to reduce the influence of high amplitude low-frequency components caused by motion artifacts and bad electrode-skin contact, as well as reducing power line-noise interferences. After filtering, stimulation sequences have been extracted and split into shorter observation windows starting at the beginning of the visual stimulation. Maximal canonical correlation has been calculated for each stimulation frequency and its second harmonic. Table 1 presents the detection accuracy and ITR for the 6-degree-of-freedom BCI-system under test in 0.5, 1, 1.5, 2, 2.5, 3, 3.5, and 4- second observation windows. Figure 3 shows the individual ITR and detection accuracy for each participant for the proposed observation windows. Detection accuracy has been proven to increase along with the observation window length, achieving a maximum of 65% in the 4-second observation window. The maximum information transfer rate, 21 bits/minute, was obtained in the 1.5-second observation window. The data shows an important amount of subject variability: while some subjects achieve an excellent detection accuracy and ITR (Subject 3 has 92% classification accuracy for a 4-second observation window and a ITR of 75 bits/minute in 1.5-second windows), others deliver poor classification results (Subject 5 has 23% detection accuracy for 4-second observation windows and a maximum ITR of 6 bits/minute).

### 2) ESN Parameterization and Stimulation Frequency Detection

ESN networks have been configured to have *N_e_ = 3* input nodes and one output node. Input nodes are fed with the filtered signals coming from O1, Oz and O2. The activation function of the network nodes is set to a hyperbolic tangent. A washout duration of 250 samples has been applied.

### RC-ESN Optimal Parameterization

The optimal number of internal units, spectral radius and input scaling has been calculated through exhaustive search. The detection accuracy calculated over 4-second observation windows and 0.5-second baseline has been assessed for every combination of internal units (from 10 to 200), spectral radius (from 0 to 1) and input scaling factor (0.001, 0.01, 0.1 and 1). In order to keep the temporal dynamics, recordings (containing 15 stimulation trials) have been split into three non-overlapping time series consisting of five consecutive stimulation/non-stimulation trials. Cross-validation using the concatenation of two-time series as training set and the remaining series as test set has been employed to evaluate every tuple under test. In each stimulation sequence under evaluation, the frequency maximizing the difference between the average ESN output in the 4- second observation window and the average ESN output in a 0.5-second baseline measured before the stimulation trial is selected as responsible of eliciting the visual evoked response. A spectral radius of 0.7, 140 internal units and 0.1 input scaling factor was found to deliver the best average classification accuracy across subjects (65.4%).

### RC-ESN SSVEP Detection

The previously calculated optimal spectral radius, internal units and input scaling are applied to SSVEP detection in 0.5, 1, 1.5, 2, 2.5, 3, 3.5, and 4-second observation windows using a baseline of 0.5 seconds. Cross-validation as described in Section 5.2.1 above has been used to assess the detection performance at each observation window. The average of three independent iterations has been calculated in order to reduce the random dynamical reservoir construction. Table 1 presents the calculated detection accuracy and ITR, considering the trial duration as the addition of the observation window and the initial baseline duration.Figure 4 presents individual ITR and detection accuracy for each participant. Unlike in canonical correlation analysis, observation window duration does not influence the detection accuracy of the RC-ESN-based detection. Similar average and individual detection performance is achieved for every observation window length under test. Average detection accuracy for every observation window is similar to the maximum accuracy obtained for CCA, in this case in 4-second windows. This fact boosts the maximum obtained ITR that reaches 49 bits/minute in 0.5-second windows.

### ESN Parameterization Influence in SSVEP detection

The spectral radius is intimately connected to the timescale of the reservoir and is a key ESN parameter. The impact of spectral radius is evaluated by setting the number of internal units to 140, the input scaling to 0.1 and computing the detection performance through cross fold validation as described in Section 5.2.1, for spectral radius values of between 0 and 1. Setting the spectral radius to zero kills the recurrence in the reservoir. Since feedback weights are used, such a zero-spectral-radius system would amount to an IIR filter. Figure 5 shows individual and average detection accuracy performance for different observation windows. The results reveal that performance significantly decreases for a zero-spectral radius system, proving that the system is actually exploiting the within-reservoir recurrence to perform the SSVEP detection. According to Figure 5, the average performance of all observation windows under test shows that average detection accuracy slightly improves along with the spectral radius up to a spectral radius value of 0.7.

The model size, defined by the number of internal units, is also a key training parameter. In general, more internal units can learn more complex dynamics, although it is very important to avoid over-fitting of the training set if a large number of internal units is used. The influence of model size in SSVEP detection performance is evaluated setting the spectral radius to 0.7, the input scaling factor to 0.1 and calculating the detection performance through three-fold cross validation as described in Section 5.2.1, for a number of internal units ranging from 10 to 200. In Figure 5 the detection performance of the model size under evaluation is presented. The results show that average detection performance increases asymptotically with model size.

**Table 1.** Detection accuracy percentage and ITR (within brackets) in bits / minute

|  | 0.5'' |  | 1'' |  | 1.5'' |  | 2'' |  | 2.5'' |  | 3'' |  | 3.5'' |  | 4'' |  |
| --- | --- | --- | --- | --- | --- | --- | --- | --- | --- | --- | --- | --- | --- | --- | --- | --- |
|  | cca | esn | cca | esn | cca | esn | cca | esn | cca | esn | cca | esn | cca | esn | cca | esn |
| S1 | 17<br>(0) | 77<br>(77) | 36<br>(9) | 80<br>(55) | 52<br>(19) | 84<br>(47) | 58<br>(18) | 84<br>(38) | 66<br>(20) | 84<br>(32) | 68<br>(18) | 86<br>(29) | 76<br>(20) | 91<br>(29) | 76<br>(18) | 86<br>(22) |
| S2 | 19<br>(0) | 40<br>(13) | 28<br>(3) | 40<br>(9) | 39<br>(8) | 42<br>(8) | 44<br>(9) | 42<br>(6) | 54<br>(12) | 42<br>(5) | 62<br>(15) | 39<br>(3) | 67<br>(15) | 40<br>(3) | 69<br>(14) | 42<br>(3) |
| S3 | 28<br>(6) | 89<br>(108) | 74<br>(70) | 93<br>(81) | 90<br>(75) | 93<br>(61) | 88<br>(51) | 94<br>(51) | 94<br>(51) | 92<br>(40) | 92<br>(40) | 90<br>(32) | 91<br>(33) | 91<br>(29) | 92<br>(30) | 90<br>(25) |
| S4 | 16<br>(0) | 44<br>(18) | 22<br>(1) | 53<br>(19) | 32<br>(4) | 53<br>(15) | 43<br>(8) | 53<br>(12) | 44<br>(7) | 57<br>(12) | 51<br>(9) | 53<br>(9) | 53<br>(8) | 54<br>(8) | 66<br>(12) | 51<br>(6) |
| S5 | 23<br>(3) | 52<br>(29) | 32<br>(6) | 49<br>(16) | 22<br>(0) | 51<br>(13) | 21<br>(0) | 49<br>(10) | 19<br>(0) | 50<br>(8) | 23<br>(0) | 48<br>(7) | 28<br>(1) | 44<br>(4) | 23<br>(0) | 44<br>(4) |
| Avg | 21<br>(2) | 61<br>(49) | 38<br>(15) | 63<br>(36) | 47<br>(18) | 64<br>(29) | 51<br>(14) | 65<br>(23) | 55<br>(15) | 65<br>(20) | 59<br>(14) | 63<br>(16) | 63<br>(13) | 64<br>(15) | 65<br>(12) | 63<br>(12) |

**Fig. 4.**
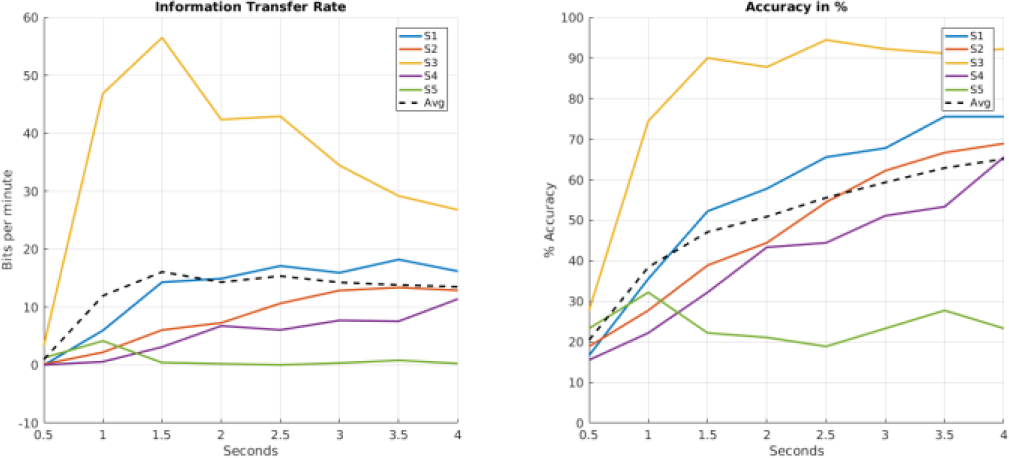
Information transfer rate (left) and detection accuracy (right) at different observation-window length using standard CCA detection.

**Fig. 5.**
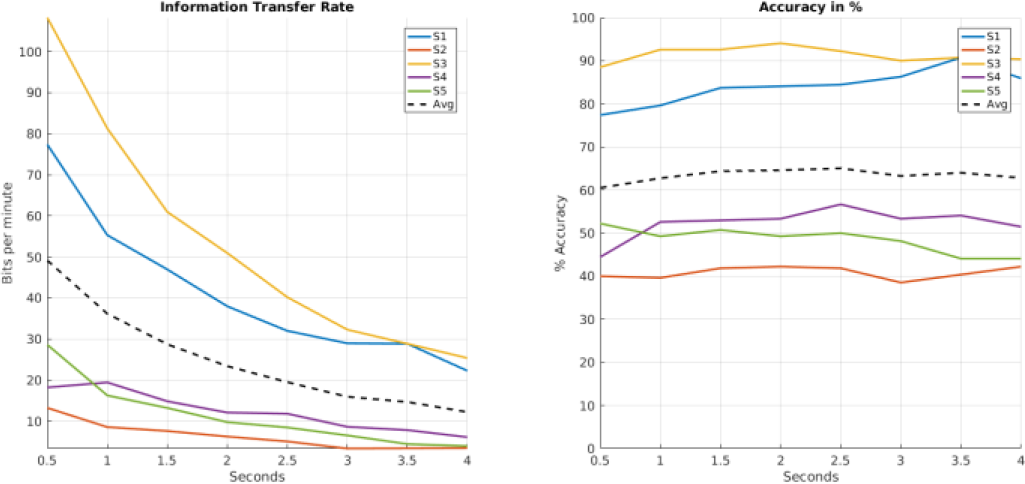
Information transfer rate (left) and detection accuracy (right) at different observation-window length using ESN-based detection.

**Fig. 6.**
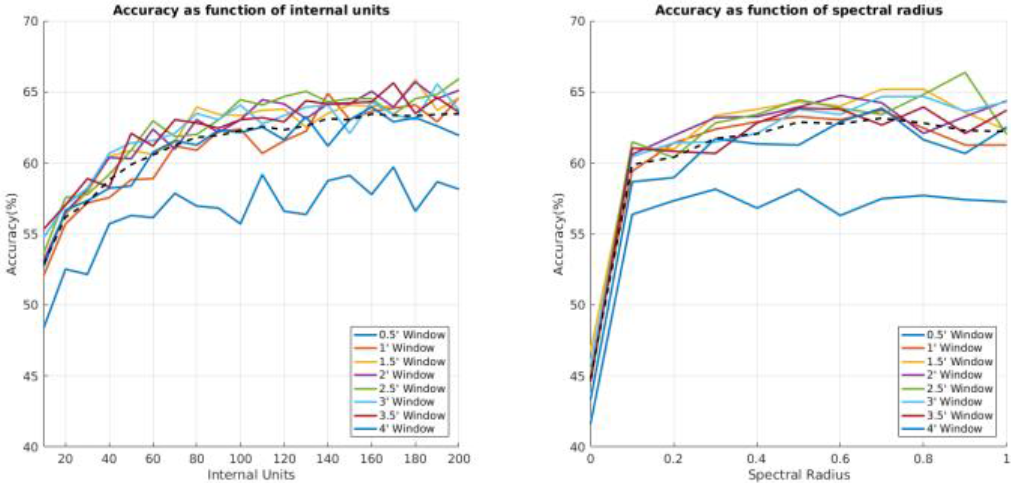
Detection accuracy of the ESN-based method as function of the number of internal units (left) and spectral radius (right).

### Individual Stimulation Frequency Detection

SSVEP is a subject-dependent phenomenon in which a given stimulation frequency has proved to range from excellent classification accuracy to random classification among subjects [36]. The feasibility of an SSVEP-based system thus strongly depends on the appropriate individualization of used stimulation frequencies. Table 2 presents the detection accuracy for each stimulation frequency obtained for the observation windows delivering largest average detection accuracy along all stimulation frequencies for CCA and ESN-based methodologies, respectively 4 and 2 seconds. ESN-based methodologies significantly improve the average detection accuracy of subjects 1 and 5, while standard CCA performs better in subject 2 and 4. Both methodologies deliver similar average classification performance for subject 3. Results show that the ESN-based method significantly improves classification in some stimulation frequencies and subjects compared to CCA (subject 1, stimulation frequency 20 Hz) and vice versa (subject 2 stimulation frequency 14 Hz). These classification differences may prove the elicitation of evoked responses of different nature, explaining why stationary and dynamical detection methodologies perform differently. The brain response to a flickering stimulation has traditionally been considered to be a steady-state system, in which the effect elicited is considered to be unchanging in time. In the cases in which reservoir computing outperforms CCA, complex dynamical patterns and/or complex synchronization among EEG channels may have been detected. The complex nature of the elicited visual stimulation response shall be studied further in order to complete the definition of steady-state visual evoked potentials. In any case, our results indicate that standard CCA and ESN-based methodologies are complementary in terms of SSVEP detection.

**Table 2.** Detection accuracy percentage of canonical correlation analysis (4-seconds observation window) and proposed ESN-based methodologies (2-seconds observation window)

|  | Subject 1 |  | Subject 2 |  | Subject 3 |  | Subject 4 |  | Subject 5 |  |
| --- | --- | --- | --- | --- | --- | --- | --- | --- | --- | --- |
|  | cca<br>w=4'' | esn<br>w=2'' | cca<br>w=4'' | esn<br>w=2'' | cca<br>w=4'' | esn<br>w=2'' | cca<br>w=4'' | esn<br>w=2'' | cca<br>w=4'' | esn<br>w=2'' |
| 12Hz | 100 | 87 | 100 | 87 | 100 | 100 | 67 | 6 | 67 | 20 |
| 14Hz | 93 | 80 | 80 | 33 | 100 | 100 | 80 | 33 | 40 | 53 |
| 16Hz | 80 | 80 | 66 | 13 | 100 | 100 | 80 | 87 | 13 | 53 |
| 18Hz | 87 | 87 | 46 | 47 | 100 | 87 | 80 | 53 | 6 | 60 |
| 20Hz | 53 | 80 | 33 | 13 | 93 | 100 | 53 | 80 | 13 | 87 |
| 22Hz | 40 | 93 | 86 | 8 | 60 | 80 | 33 | 67 | 0 | 33 |
| Avg | 75 | 84 | 68 | 39 | 92 | 91 | 65 | 50 | 23 | 50 |

## VI. Discussion and Conclusions

We have presented a reliable SSVEP-detection methodology based on reservoir computing with online capabilities. Its performance has been successfully compared to standard canonical correlation analysis for the construction of a BCI system of six degrees of freedom. The information transfer rate of the hereby proposed approach does not overcome state of the art high-speed SSVEP approaches such as Nakanishi’s [39]. However it is limited in terms of degrees of freedom (6 instead of 32), covers a larger stimulation frequency range (from 6 to 22 instead of from 8 to 15) without phase-coding and uses a smaller number of electrodes.

The performance of our proposed ESN-based method proved to be non-dependent from observation window length, delivering similar detection accuracy for windows ranging from 0.5 to 4 seconds (from 61% to 65%). In contrast, CCA showed a strong dependence on the stimulation window duration, with 0.5-second observation windows delivering random classification, and accuracy increasing with window length, reaching a maximum detection accuracy of 65% in 4- second windows. These results highlight the communication capabilities of the ESN-based method, which achieves an average information transfer rate of 49 bits/minute (with a maximum ITR of 108 bits/minute obtained for a single subject), compared with the maximum average information transfer rate of 21 bits/minute achieved by the CCA method. The ESN-based method achieves an excellent information transfer rate for a six-degrees-of-freedom BCI system compared to other state of the art high-rate SSVEP systems [2,37,38].

SSVEP is a subject-dependent technique in which the stimulation frequency defines the classification performance. In this study, a wide range of frequencies ranging from 10 to 22 Hz in steps of 2 Hz is used to simulate a six-degrees-of-freedom SSVEP-based BCI system. Individualized stimulation frequency selection is expected to improve the detection accuracy of both CCA and ESN-based methods. Both methodologies have proved to be complementary in terms of detection accuracy. The ESN and CCA-based methods proved to deliver excellent detection accuracy at different stimulation frequencies. For instance, in our study Subject 5 offered 87% detection accuracy at 20Hz stimulation when using the ESN-based detection methodologies, and near-to-random classification with the CCA-based method. In contrast, Subject 2 showed 80% detection accuracy for CCA-based methods and 33% for the ESN-based technique. A possible explanation for this behavior could be that in different subjects each stimulation frequency may elicit evoked responses of different nature, in terms of their dynamical/stationary characteristics. Specifically, reservoir computing methods are able to detect complex dynamical patterns and/or complex synchronization among EEG channels, in contrast to stationary patterns detected by canonical correlation analysis. The brain response to a flickering stimulation has traditionally been considered to be a steady-state system, in which the effect elicited is considered to be unchanging in time. The hypothesis of complex dynamical activity elicited by visual repetitive stimulation shall be further studied in order to complete, if confirmed, the definition of steady-state visual evoked response.

## Acknowledgment

We want to thank all members of the AsTeRICS project consortium; special acknowledgements go to the AsTeRICS users who helped us to test and improve the system. We also kindly thank the developers of the Oger toolbox [40].

